# Non-invasive vagus nerve stimulation boosts mood recovery after effort exertion

**DOI:** 10.1101/2020.07.21.214353

**Authors:** Magdalena Ferstl, Vanessa Teckentrup, Wy Ming Lin, Franziska Kräutlein, Anne Kühnel, Johannes Klaus, Martin Walter, Nils B. Kroemer

## Abstract

Mood plays an important role in our life which is illustrated by the disruptive impact of aberrant mood states in depression. Although vagus nerve stimulation (VNS) has been shown to improve symptoms of depression, the exact mechanism is still elusive, and it is an open question whether non-invasive VNS could be used to swiftly and robustly improve mood. Here, we investigated the effect of left- and right-sided transcutaneous auricular VNS (taVNS) versus a sham control condition on mood after exertion of physical and cognitive effort in 82 healthy participants (randomized cross-over design). Using linear mixed-effects and hierarchical Bayesian analyses of mood ratings, we found that 90 min of either left-sided or right-sided taVNS improved positive mood (*b* = 5.11, 95% credible interval, CI [1.39, 9.01], 9.6% improvement relative to the mood intercept, BF_10_= 7.69, *p*_LME_ = .017), yet only during the post stimulation phase. Moreover, lower baseline scores of positive mood were associated with greater taVNS-induced improvements in motivation (*r* = −.42, 95% CI [−.58, −.21], BF_10_ = 249). We conclude that taVNS boosts mood after a prolonged period of effort exertion with concurrent stimulation and that acute motivational effects of taVNS are partly dependent on initial mood states. Collectively, our results show that taVNS may help quickly improve affect after a mood challenge, potentially by modulating interoceptive signals contributing to reappraisal of effortful behavior. This suggests that taVNS could be a useful add-on to current behavioral therapies.

## Introduction

We are often faced with demanding cognitive and physical tasks. Completing effortful tasks can leave us feeling drained and exhausted, leading to a dampened mood (Broderick, 2005; Erber & Erber, 1994; Erber & Tesser, 1992; Kron et al., 2010). Imagine that you need to work through a pile of papers by the end of the day. Despite your positive mood in the morning, you will probably feel exhausted afterwards and it might take long to return to the positive feeling you had before. Accordingly, we experience a wide range of mood states that encompasses a spectrum from positive to negative moods every day (Clark & Watson, 1988; R. E. Thayer, 1997). In contrast to temporary alterations in mood, persistent negative mood that is present in affective disorders incurs a debilitating individual burden (Ferrari et al., 2013) and high socio-economic costs (König et al., 2019). Thus, it is crucial to identify potential causes of mood alterations and to investigate ways to alleviate pathologically relevant mood disturbances.

As illustrated by going to work and completing assignments, one factor that affects our moods are cognitive and physical tasks that require effort to complete. On the one hand, exerting cognitive effort after positive and negative mood induction leads to a dampening of the induced mood (Broderick, 2005; Erber & Erber, 1994; Erber & Tesser, 1992; Kron et al., 2010). Additional mood regulation processes have been observed to mitigate induced negative mood states with a concurrent increase in positive mood (Erber & Erber, 2000; Forkosh & Drake, 2017; Kim & Kanfer, 2009; Sirois, 2014). On the other hand, exerting physical effort may increase or decrease mood depending on how much effort was exerted. Higher training intensities can lead to an increase in negative mood immediately following training (Morgan et al., 1987; Oliveira et al., 2013; Selmi et al., 2018), in part due to unpleasant interoceptive cues for high exercise intensity (Ekkekakis, 2009). Nevertheless, effort exertion may lead to long-term improvements in mood and symptoms of depression, demonstrating the relevance of post-acute recovery (Brosse et al., 2002; Schuch et al., 2016; Stathopoulou et al., 2006). Taken together, these findings point to task exertion as a core modulator of affect, potentially leading to aberrant mood states if mood does not recover quickly after exertion.

Since prolonged negative mood is a key symptom of major depression and related mental disorders ((*Diagnostic and Statistical Manual of Mental Disorders: DSM-5*, 2013) and ICD-10 (World Health Organization, 2004)), treatments of affective disorders often target improvements in mood (Sin & Lyubomirsky, 2009). Current treatment for depression is heterogeneous and can include medication, behavioral therapies, or brain stimulation techniques such as deep brain stimulation (Delaloye & Holtzheimer, 2014) or vagus nerve stimulation (Cimpianu et al., 2017; Howland, 2014). When VNS is applied invasively, the device is usually implanted in the chest wall with wires stimulating the vagus nerve at the neck (George et al., 2000). Previous studies have reported improvements in mood states and depressive symptoms after VNS (Elger et al., 2000; George et al., 2005, 2008; Rush et al., 2005; Schlaepfer et al., 2008) which increased with the duration of use (Berry et al., 2013). Given the invasive nature of VNS, a non-invasive alternative is needed as it would greatly expand its potential use as an adjunct treatment (Ventureyra, 2000). Indeed, transcutaneous auricular vagus nerve stimulation (taVNS) was developed as a non-invasive variant of VNS where the vagus nerve is stimulated through the skin of the auricle (Fallgatter et al., 2003). Preliminary studies have shown that non-invasive taVNS improves mood (Kraus et al., 2007) and depressive symptoms (Liu et al., 2016; Tu et al., 2018; Wu et al., 2018) similar to invasive VNS. Such improvements could be due to alterations in brain activity in the prefrontal cortex and the limbic system after invasive (Nahas et al., 2007; Pardo et al., 2008) and non-invasive VNS (Badran et al., 2018; Yakunina et al., 2017). Since these mesocorticolimbic structures are characteristically dysregulated in depression (Anand et al., 2005; Groenewold et al., 2013; Iseger et al., 2020) and critically involved in cost-benefit decision-making (Husain & Roiser, 2018; Kroemer et al., 2014, 2016), taVNS may provide a means to improve mood, particularly after effort exertion.

Despite emerging evidence for beneficial effects of taVNS on mood especially when used chronically (Hein et al., 2013; Rong et al., 2016), studies on the acute effects of taVNS on mood remain surprisingly scarce (Kraus et al., 2007). Moreover, none of the studies include a challenge that temporarily reduces mood. To close this gap, we investigated the effects of acute taVNS versus sham on mood after the exertion of effort and after a mood recovery period. To measure mood states, we used the Positive and Negative Affect Schedule (Watson et al., 1988) before, during, and after stimulation. Based on previous results, we hypothesized that taVNS enhances positive mood and attenuates negative mood compared to sham stimulation. In line with the prediction, we found marked improvements of positive mood in the post stimulation phase but not directly after effort exertion during taVNS. Crucially, the hierarchical Bayesian modeling results provide strong evidence for a possible benefit of taVNS on mood during a post stimulation recovery period and may be of particular interest for studies investigating applications of acute taVNS in mood disorders.

## Methods

### Participants

Eighty-five volunteers participated in the study. To be included in the analysis, each participant had to complete two sessions: one after stimulation of the cymba conchae and one after a sham stimulation at the earlobe. Thus, we had to exclude 3 participants who did not finish the second experimental session, leading to a sample size of *N* = 82 with complete mood ratings. As reported before (Neuser et al., 2020), one additional participant had to be excluded from the correlation analysis involving taVNS-induced changes in motivation because of an incorrectly assigned maximum of the button press frequency for the effort task precluding comparisons of the two sessions. Out of the 82 participants, 42 completed the study during left-sided taVNS, whereas 40 completed it during right-sided taVNS. All participants went through a telephone screening to ensure that they were healthy, spoke German, and were right-handed (47 women; *M*_Age_= 24.6 years ± 3.5; *M*_BMI_= 23.1 kg/m^2^ ± 3.01; 17.9 - 30.9). The study was approved by the local ethics committee and was conducted in accordance with the ethical code of the World Medical Association (Declaration of Helsinki). Participants provided written informed consent at the beginning of Session 1 and received either monetary compensation (32€ fixed amount) or course credit for their participation after completing the second session. Moreover, they received money and breakfast depending on their performance during the tasks.

### Experimental procedure

Experimental sessions were conducted in a randomized, single-blind crossover design (Fig. 1). Participants were required to fast overnight. Each session began in the morning and lasted about 2.5h. After measuring physiological parameters such as pulse and weight as well as assessing information about last food intake, the participants were instructed that they would be collecting energy and money points throughout the session depending on their performance in the following tasks. Earned energy points would be converted into participants’ breakfast consisting of cereal and milk, scaled according to the number of points they won. Water was provided ad libitum during the experiment.

**Figure 1:**
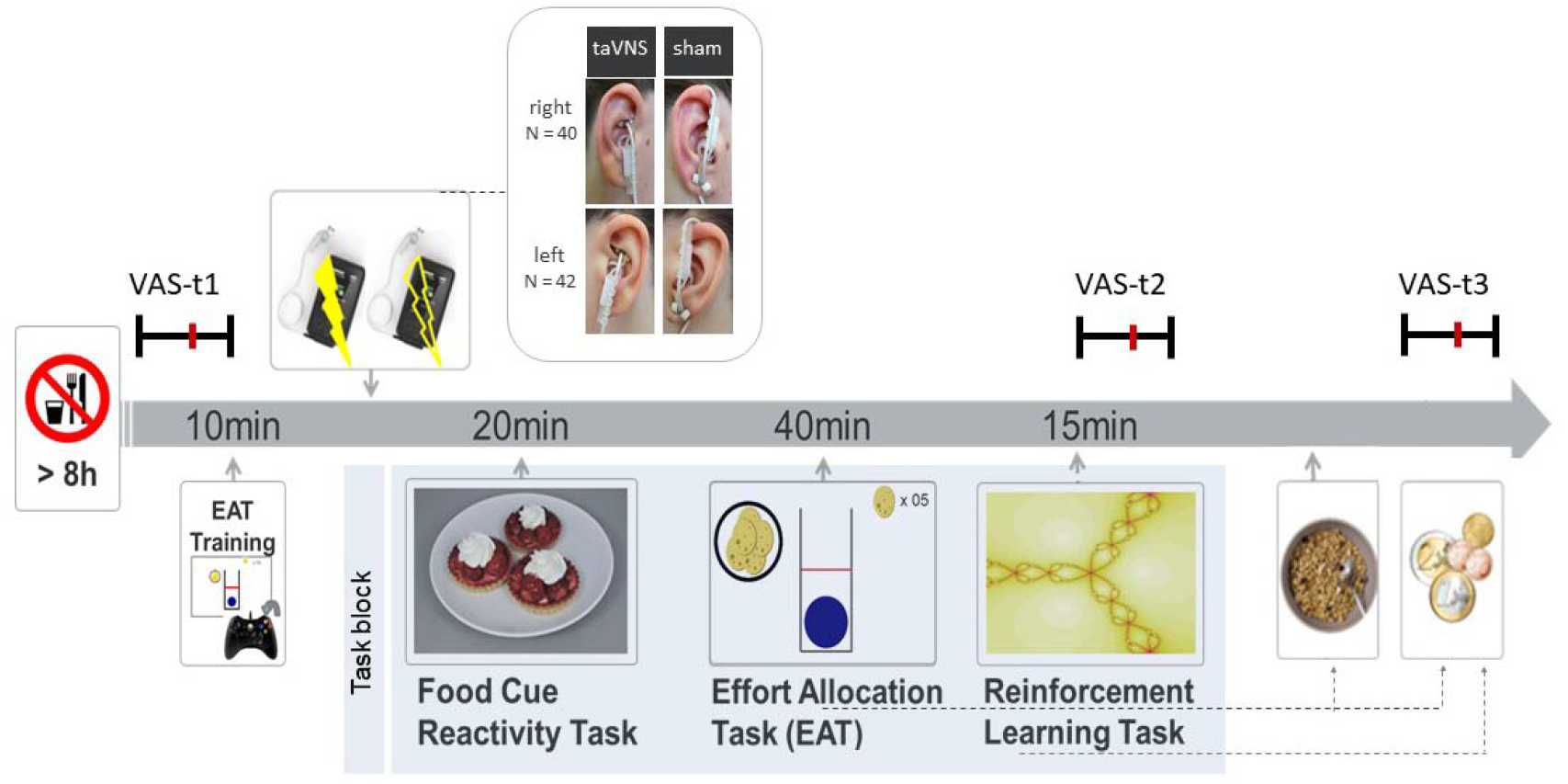
Experimental procedure and schematic summary of the task block. The timeline shows the experimental procedure with stimulation beginning before a block of three tasks. State measurements were taken before stimulation, during stimulation as well as after the tasks, and after stimulation using visual analog scales (VAS). The food cue reactivity task required quick ratings of wanting and liking. The effort allocation task required physical effort to obtain rewards by pressing a button vigorously. The reinforcement learning task required cognitive effort to track which actions following a cue maximize reward and minimize punishments.

Next, participants responded to state questions based on the PANAS (Watson et al., 1988) using visual analog scales (VAS). A practice session of the effort task followed where the maximum frequency of button press was recorded to adjust the task to each participant’s individual ability.

Afterwards, the taVNS electrode was placed either on the left (*N* = 42) or the right (*N* = 40) ear. Strips of surgical tape secured the electrode and the cable in place. For every session and stimulation condition, the stimulation strength was individually assessed using VAS ratings of pain to match the subjective experience. Stimulation was initiated at an amplitude of 0.1 mA and increased by the experimenter by 0.1-0.2 mA at a time. Participants rated the sensation after every increment until ratings corresponded to a “mild pricking”. The stimulation remained active at this level.

Participants then completed a food-cue reactivity task (~20 min) as well as the effort task (~40 min), which tested the willingness to expend physical effort in return for rewards. Since the default stimulation protocol of the NEMOS taVNS device alternates between 30 s of on and off phases, the experimenter manually switched on the stimulation phases of the device at the onset of the effort phases of the task. Moreover, participants completed a reinforcement learning task (Kühnel et al., 2020), which required cognitive effort (~15 min).

After completing the task block, participants entered state VAS ratings for the second time. Then, participants had the taVNS electrode removed and received their breakfast and a snack according to the energy points earned in the effort allocation task. They were instructed that this was their food reward and that they could eat as much as they liked. Next, participants completed state VAS ratings for a third time. To conclude the first session, all participants received their monetary winnings earned in the effort allocation task and reinforcement learning task as part of the compensation. Both visits were conducted at approximately the same time within a week, usually within 3-4 days, and followed the same standardized protocol (for details on the procedure, see (Neuser et al., 2020).

### Mood ratings

Participants responded to state questions at three time points: before starting the tasks and stimulation (baseline VAS), after completing the task block during the stimulation (stimulation VAS), and finally about 20 min after completing the tasks and the stimulation period and after receiving the earned rewards including breakfast (post stimulation VAS). The questionnaire was presented on a computer screen as VAS using the joystick on an Xbox 360 controller (Microsoft Corporation, Redmond, WA). For assessing the participants’ mood, we used the PANAS (Watson et al., 1988). Positive affect can be described as a state of high energy and enthusiasm, whereas negative affect comprises different aversive feelings such as anger, guilt, or nervousness.

### Effort allocation task

In the effort allocation task, participants had to collect food and money tokens throughout the task by exerting effort (i.e., repeatedly pressing the right trigger button with the right index finger). The task was adapted from (Meyniel et al., 2013) and used frequency of button presses instead of grip force to measure physical effort (Neuser et al., 2020). Trials varied in the type of reward, which was always indicated by money or food symbols on the screen. Moreover, we varied the magnitude of the prospective reward (low vs. high) as well as the difficulty (easy vs. hard) to obtain the rewards. After every effort phase of a trial, participants were presented sequentially with two VAS questions inquiring about the exertion for and the wanting of the reward at stake.

### taVNS device

To stimulate the auricular branch of the vagus nerve, we used the NEMOS® stimulation device (cerbomed GmbH, Erlangen, Germany). These devices have been previously used in clinical trials (Bauer et al., 2016; Kreuzer et al., 2012) and proof-of-principle neuroimaging studies (Frangos et al., 2015). The stimulation protocol for the NEMOS is preset to a biphasic impulse frequency of 25 Hz with a stimulation duration of 30 s, followed by a 30 s off phase. The electrical current is transmitted by a titanium electrode placed at the cymba conchae (taVNS) or earlobe (sham) of the ear (Frangos et al., 2015). To match the subjective experience of the stimulation, intensity was determined for each participant and each condition individually to correspond to mild pricking (Kühnel et al., 2020; Neuser et al., 2020; Teckentrup et al., 2020).

### Data analysis

#### Statistical modeling of mood ratings

Although VAS scales are commonly used to measure mood, there are several potential pitfalls that have to be addressed in the analysis (Heller et al., 2016; Vautier, 2011). For example, change scores after an intervention are often strongly correlated with baseline levels and this statistical dependence must be modeled accordingly (Vickers & Altman, 2001). Therefore, we used linear mixed-effects as well as hierarchical Bayesian analyses to analyze taVNS-induced changes in mood based on all collected ratings. Compared to ANOVAs, linear mixed-effects analyses are advantageous because they enable the estimation of “fixed” effects of the stimulation and other design elements (“immutable” differences that are shared across the group), “random” effects (persistent differences of individuals from a group average), and noise (non-reproducible differences due to measurement error). A repeated measures ANOVA can be regarded as a simple and restricted mixed-effects model (Raudenbush & Bryk, 2002). Compared to mixed-effects analyses, hierarchical Bayesian models are advantageous because they provide several crucial extensions (Gelman & Hill, 2006). First, it is possible to use informed priors to improve the estimation and precision of a model (Bürkner, 2017; Gelman & Hill, 2006). For example, a useful constraint is to restrict the range of the VAS ratings from 0 to 100. Second, it is possible to sample the posterior distribution of a parameter (Bürkner, 2017). This posterior distribution represents the updated beliefs about the parameters after having seen the data in terms of a probability distribution (Gelman & Hill, 2006). In contrast, linear mixed-effects analyses estimate differences in parameters compared to null hypotheses based on distributional statistics of the data (Raudenbush & Bryk, 2002). Third, by sampling the posterior distribution, we can estimate the likelihood of the alternative hypothesis, including the likelihood of a desired effect size such as taVNS-induced changes in mood, instead of estimating the probability of observing a change of a given magnitude if the null hypothesis were true.

Therefore, in addition to frequentist *p*-values estimated by the linear mixed-effects model, we also estimated a corresponding hierarchical Bayesian model. Model 1+ predicted the outcome positive mood ratings of all individual items using the factors run (1: baseline VAS, 2: stimulation VAS, 3: post stimulation VAS) and stimulation as well as their interaction (random slopes and intercept). Model 1-predicted the outcome negative mood ratings of all individual items using the same set of predictors (for prior settings and R code used to fit the models, see SI).

#### Estimation of invigoration and maintenance of effort as a motivational index

For isolating invigoration and maintenance of effort as motivational indices, we segmented the behavioral data into work and rest segments as reported in a previous publication (Neuser et al., 2020). Invigoration of effort was computed by estimating the slope of the transition between the relative frequency of button presses during a rest segment and their initial plateau during the subsequent work segment based on the MATLAB findpeaks function. Maintenance of effort was computed as the average relative frequency of button press over a given trial, which is equivalent to the area under the curve. As previously reported (Neuser et al., 2020), taVNS effects were calculated using two univariate mixed-effects models. Briefly, either invigoration or maintenance of effort were predicted based on stimulation (taVNS, sham), reward type (food, money), reward magnitude (low, high), difficulty (easy, hard; all dummy coded), the interaction between reward magnitude and difficulty as well as interactions between stimulation and all other terms. At the participant level, stimulation order and stimulation side (mean centered) were included. To account for deviations from fixed group effects, random slopes and intercepts were modeled for all predictors.

### Statistical threshold and software

For our analyses, we used a two-tailed α ≤ .05. Mixed-effects analyses were conducted with HLM v7 (effort invigoration and maintenance; Raudenbush et al., 2011) and lmerTest in R (positive and negative mood, Kuznetsova et al., 2017).

Hierarchical Bayesian analyses were conducted with brms in R (Bürkner, 2017). To determine the evidence for the alternative hypothesis (i.e., taVNS improves mood) provided by our results, we calculated corresponding Bayes Factors (BF). For the correlation with taVNS-induced changes in motivation, we used a stretched beta prior = 0.7 (to reflect that large correlations are less likely a priori) as implemented in JASP v0.9 (JASP Team, 2019). We processed data with MATLAB vR2019a and SPSS v26 and plotted results with R v3.6.0 (R Core Team, 2019), including bayesplot (Gabry & Mahr, 2019) and JASP.

## Results

### taVNS increases positive mood after the stimulation phase

To estimate taVNS-induced changes in mood, we first used a linear mixed-effects model predicting positive or negative mood as outcomes based on the factors run and stimulation as well as their interaction. To account for inter-individual differences in taVNS-induced changes, the intercept and slopes were modeled as random effects. As expected, the demanding tasks conducted during the stimulation phase led to a decrease in positive mood (fixed effect of stimulation VAS, Run 2: *b* = −7.50, *t* = 4.81, *p* < .001) that had only partly recovered 20-min after the stimulation (fixed effect of post stimulation VAS, Run 3: *b* = −3.79, *t* = 2.51, *p* = .014). However, there was no increase in negative mood during the stimulation phase (fixed effect of stimulation VAS, Run 2: *b* = 0.77, *t* = 0.79, *p* = .43) and it even improved slightly 20-min after the stimulation (fixed effect of post stimulation VAS, Run 3: *b* = −1.93, *t* = 2.27, *p* = .026).

In line with the randomization of stimulation conditions, there was no main effect of taVNS on positive and negative mood at baseline (*p*s > .48). Crucially, we observed a significant taVNS-induced improvement of positive mood in the post stimulation VAS (interaction Run 3 x Stimulation: *b* = 4.30, *t* = 2.44, *p* = .017, Fig. 2b), but not in the stimulation VAS (interaction Run 2 × Stimulation: *b* = 1.12, *t* = 0.63, *p* = .53). Likewise, we observed an taVNS-induced improvement of negative mood in the post stimulation VAS, but this effect was not significant, likely due to the restricted range of negative mood ratings in healthy participants (interaction Run 3 × Stimulation: *b* = −1.25, *t* = 1.05, *p* = .30, Fig. 2a).

**Figure 2:**
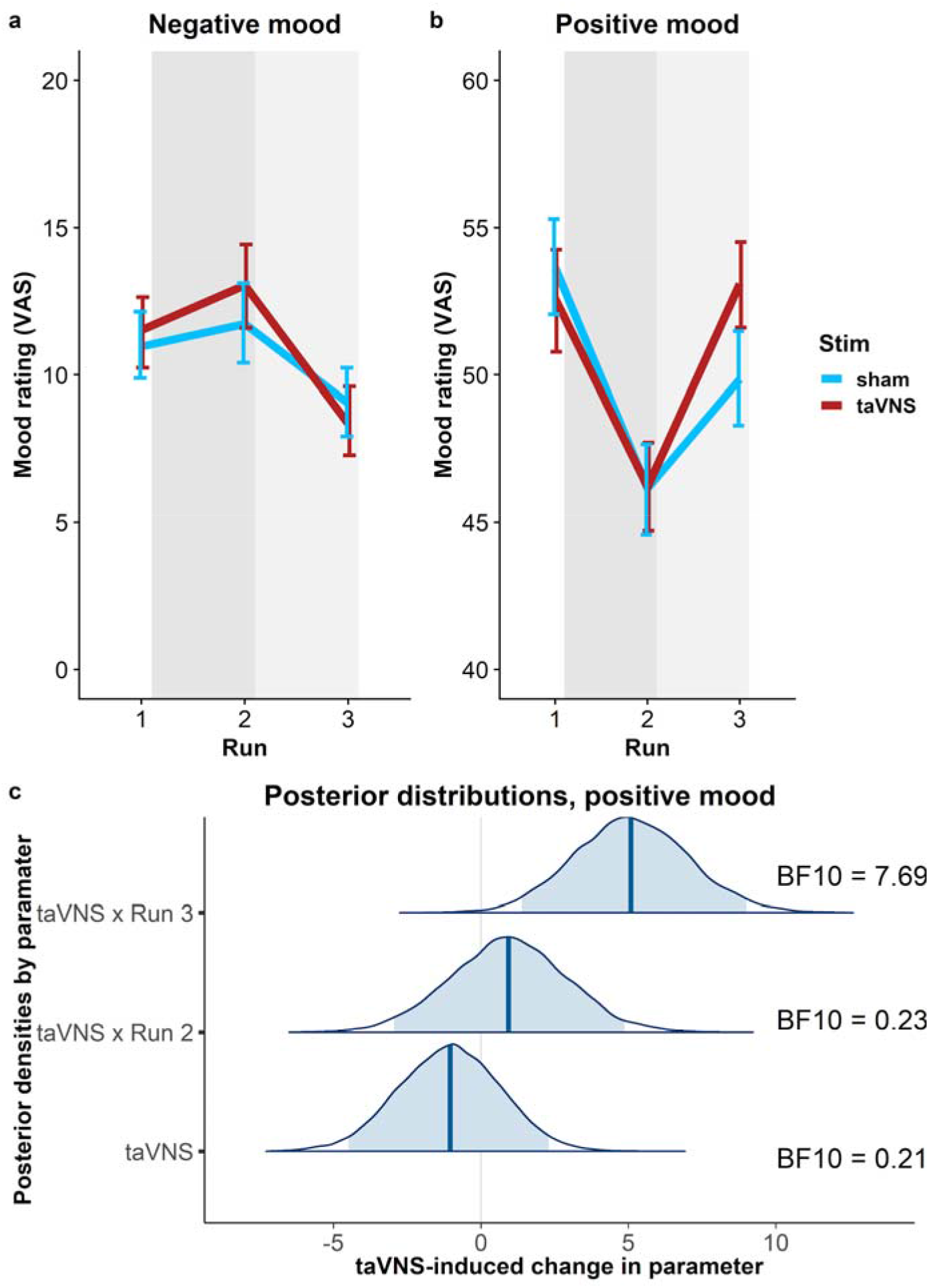
Transcutaneous auricular vagus nerve stimulation (taVNS) increases positive mood in the post stimulation visual analog scales (Run 3). A: No significant changes in negative mood. B: Significant increase in positive mood 20 min post stimulation (light gray shading), but not during stimulation (gray shading). Error bars denote 95% confidence intervals derived via bootstrapping. C: Posterior distribution of the taVNS-induced change in positive mood (95% credible interval in light blue, median in dark blue). BF_10_ = Bayes factor for the alternative hypothesis

The mixed-effects results were corroborated by the hierarchical Bayesian analysis. Crucially, the Bayesian model provided a higher estimate of the taVNS-induced increase in positive mood with a credible interval that clearly exceeded zero (interaction Run 3 × Stimulation: *b* = 5.11, *e* = 1.95, 95% CI [1.39, 9.01], Fig. 2c). Consequently, the BF provided strong support for a taVNS-induced increase in the post stimulation VAS (BF_10_ = 7.69), while there was strong support for the null hypothesis at baseline (BF_10_ = 0.21) and in the stimulation VAS (BF_10_ = 0.23). Moreover, contrasting taVNS-induced changes in the stimulation VAS versus the post stimulation VAS revealed a conclusive posterior probability, P(H1_R3_>H1_R2_|D) = .99, that the effect was larger during mood recovery after the stimulation had ended. In contrast, due to the restricted range of negative mood ratings in many participants, the hierarchical Bayesian model failed to converge.

### Baseline mood moderates taVNS-induced improvements in motivation

Since the exertion of effort during the task block had a strong effect on mood, we assessed whether taVNS-induced changes in mood and effort were associated.

We have previously shown that effort exerted during the task can be dissociated into invigoration and maintenance of effort, and taVNS only increased invigoration (Neuser et al., 2020). Notably, lower baseline scores of positive mood were associated with stronger taVNS-induced increases in invigoration, *r* = −.42, 95% CI [−.57, −.21], BF_10_ = 249 (Fig. 3), but not maintenance of effort, *r* = −.11, 95% CI [−.31,.11], BF_10_ = 0.27. However, taVNS-induced changes in mood in the stimulation VAS as well as the post stimulation VAS were not significantly correlated with changes in invigoration or maintenance of effort (*r*s < .2, BF_10_ < 0.4). This suggests that initial mood states moderate subsequent improvements in motivation while taVNS-induced improvements in affect and motivation are only weakly associated.

**Figure 3:**
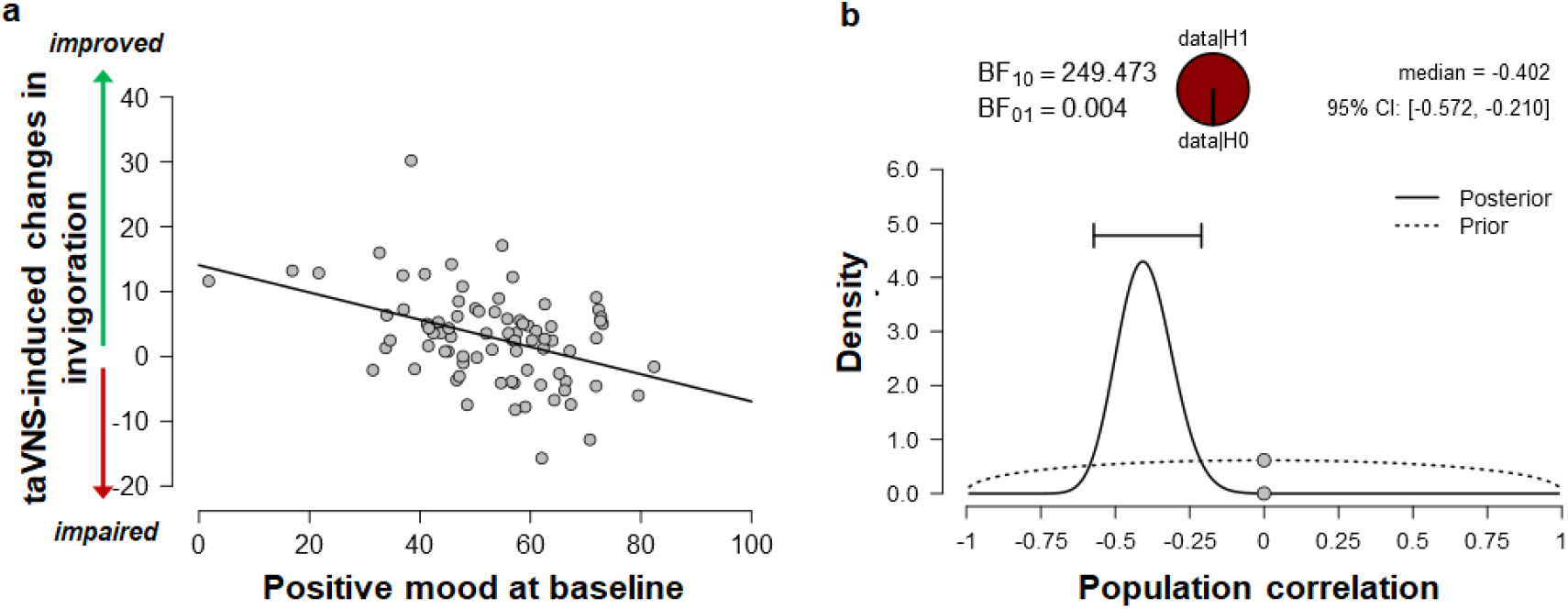
Lower positive mood at baseline is associated with greater improvements in invigoration induced by transcutaneous auricular vagus nerve stimulation. A: Scatter plot of positive mood ratings at baseline and taVNS-induced changes in invigoration during the acute stimulation phase. B: Evidence for the alternative and the null hypothesis given the observed association. BF_10_ = Bayes factor for the alternative hypothesis

### taVNS-induced improvements in mood are independent of stimulation side

To assess whether the reported results are robust across stimulation sides (left or right), sex, or order, we ran control analyses in the sample with complete data. Bayesian two-sample t-tests showed unanimous support for the null hypothesis that there are no differences in mood or changes in mood due to the stimulation side (Fig. 4a), sex, or order (BF_10_ ≤ 0.66, *p*s ≥ .12). Moreover, controlling for these three variables increased the association between baseline mood scores and taVNS-induced invigoration of effort (β_corrected_ = −.49 vs. β_uncorrected_ = −.42). These results suggest that the reported mood effects are independent of the stimulation side and robust across the potential confounds order and sex. Lastly, we explored whether taVNS-induced effects are evident across positive VAS items or driven by a more specific change. A visual inspection of the single-item residuals indicated only weak specificity suggesting that the changes in mood are best explained by a general change in positive mood (Fig. 4b).

**Figure 4:**
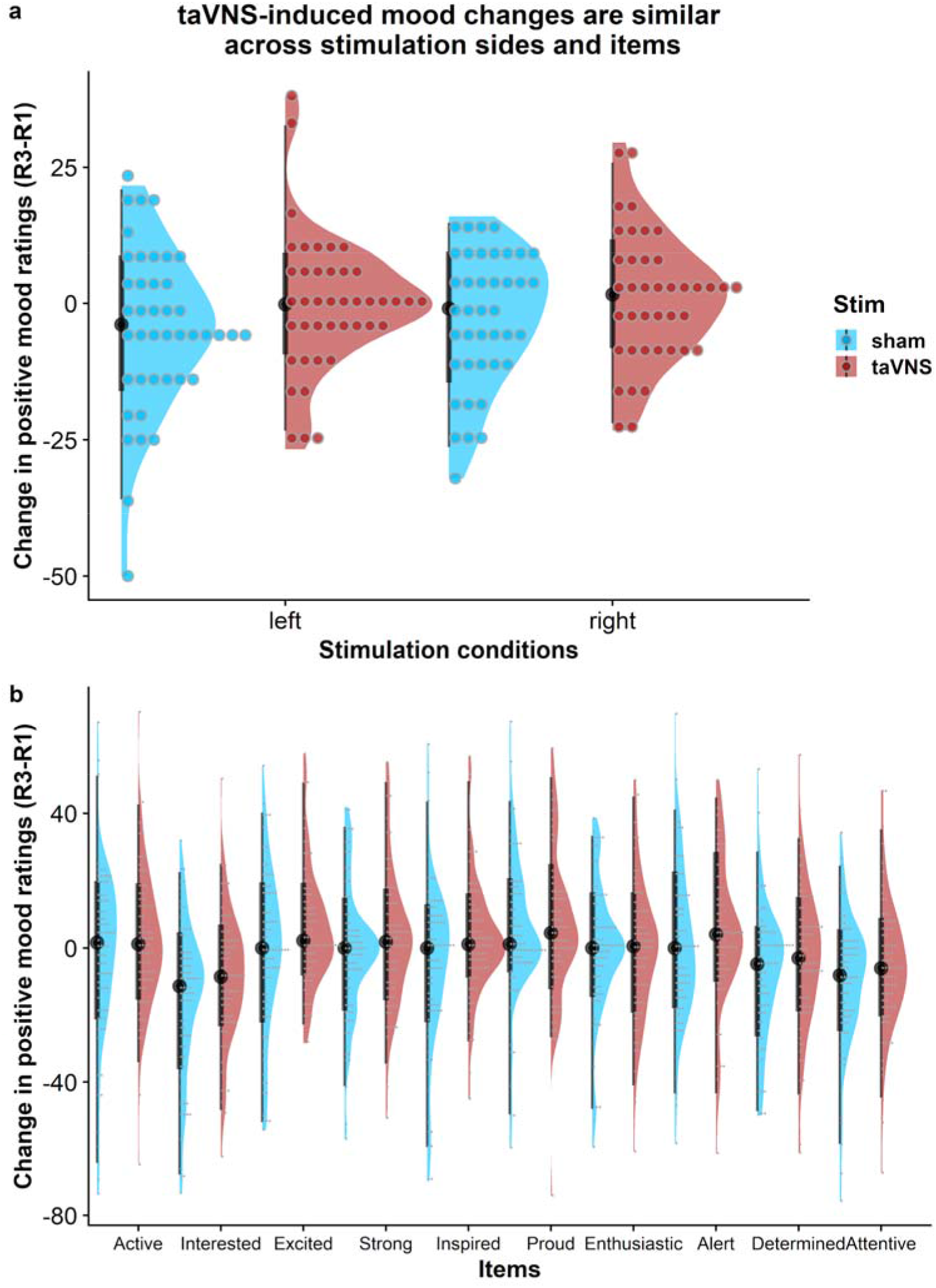
Changes in positive mood ratings during the post stimulation effects are invariant across stimulation sides and items. A: Changes in positive mood relative to baseline are depicted for each participant (dots) and as group density. Changes are comparable for transcutaneous auricular vagus nerve stimulation (taVNS) applied to the left and the right side. B: Changes in positive mood relative to baseline are largely invariant across items indicating a general effect of taVNS on mood.

## Discussion

The vagus nerve plays a vital role in the regulation of affect by providing interoceptive feedback. Whereas effects of chronic VNS on mood have been often reported (Albert et al., 2015; Elger et al., 2000; Schlaepfer et al., 2008), effects of acute non-invasive taVNS on mood have been scarcely investigated, particularly after a mood challenge. Here, we used an effortful task block to perturb mood and concurrently applied taVNS for 90 min. Using linear mixed-effects and hierarchical Bayesian models, we found that taVNS facilitated mood recovery in the post stimulation VAS, but not in the stimulation VAS directly after the task block. Furthermore, we showed that lower positive baseline mood correlated with taVNS-induced increases in invigoration of effort (Neuser et al., 2020). Crucially, taVNS-induced effects on mood did not depend on the side of stimulation, sex of the participants, or order of the sessions. Collectively, our study provides strong evidence for post-acute effects of taVNS on mood recovery after an effort challenge as well as for a moderating role of affect on motivational effects elicited by acute taVNS during an effort challenge. We conclude that taVNS could be a promising tool to improve mood regulation in response to challenges which might indicate its potential as an adjunct therapy.

Non-invasive stimulation of the vagus nerve boosted positive mood in the post stimulation VAS, approximately 20 min after stimulation and completion of effortful tasks. The facilitating effect on mood is in line with previously reported mood-enhancing effects of VNS (Howland, 2014). However, we identified a crucial temporal dissociation in acute effects of taVNS that was previously unknown. While positive mood was considerably reduced directly after the tasks (Broderick, 2005; Erber & Erber, 1994; Erber & Tesser, 1992; Kron et al., 2010) this drop in positive mood was not attenuated by taVNS. In contrast, we observed strong evidence for a faster mood recovery from an effort-induced reduction in positive affect after taVNS. A potential explanation that taVNS primarily facilitates mood recovery might be a reduced stress response induced by the tasks, as the vagus nerve has been consistently shown to play an important inhibitory role in regulating allostatic systems (Balzarotti et al., 2017; Crowley et al., 2011; Smeets, 2010; J. F. Thayer & Sternberg, 2006; Weber et al., 2010). For example, low vagal tone has been associated with an impaired recovery from stress (Porges, 2007; Weber et al., 2010) and taVNS-induced increases in vagal tone may facilitate recovery from task exertion. Notably, impaired or delayed recovery from disturbances of homeostasis have been proposed as critical factors in the development of mood disorders, suggesting that a faster recovery from stress is beneficial (Brosschot et al., 2005; McEwen, 1998). Alternatively, taVNS may increase the relative contribution of interoceptive signals forwarded via the NTS in inference processes that play a role in regulating affect (Allen et al., 2019). Put simply, taVNS may speed up recovery by facilitating the subjective inference that one’s body is in an improved state after a straining episode. Another possibility is that taVNS changes the retrospective evaluation of preceding effort for rewards once they have been received and experienced during a consummatory phase due to a modulatory effect on nigrostriatal circuits involved in cost/benefit decision-making (de Araujo et al., 2020; Neuser et al., 2020). To summarize, a taVNS-induced increase in vagal tone might alter the subjective evaluation of preceding effort during a post stimulation phase by modulating interoceptive feedback, which may ultimately facilitate mood recovery (Porges, 1992).

We further found that baseline levels of positive mood moderated taVNS-induced effects on motivation in the effort task (Neuser et al., 2020). More specifically, when positive mood was low at baseline, taVNS showed a significantly stronger impact on invigoration. A comparable dependence of taVNS effects on baseline positive mood state has been shown by Steenbergen et al., (2020) on delay discounting. A possible explanation for a modulatory role of baseline mood is that people with low baseline mood are less likely to participate in effortful, potentially mood-worsening tasks (Taquet et al., 2020), whereas elated mood states typically facilitate motivation (Young & Nusslock, 2016). Therefore, the potential of taVNS to improve motivation could be strongest for low baseline levels of positive mood. Alternatively, as previously hypothesized (Steenbergen et al., 2020), low baseline mood could be indicative of a low task-relevant arousal state which may improve more strongly by taVNS. In conclusion, taVNS may increase the motivation to engage in effort particularly in a dampened initial mood state by encouraging participants to work harder for rewards even of a lower value. In turn, this effect may help quickly restore mood once participants can reap the fruit of their labor.

Despite the conclusive evidence for improvements in mood during the post stimulation phase, there are limitations of our study that should be addressed in future research. First, we found no effects on negative mood. This is likely due to the restricted range of baseline negative mood as well as a lack of increase in negative mood in response to the tasks in our sample as the hierarchical Bayesian model failed to converge with random effects for participants. Thus, to better resolve effects on negative mood, this study should be extended either by including a more aversive task block to induce increases in negative mood or conducting it in a sample of patients with affective disorders such as major depression as negative mood is a key symptom of the disorder (*Diagnostic and Statistical Manual of Mental Disorders◻: DSM-5*, 2013). Second, although we assessed mood at three time points providing a more fine-grained resolution compared to previous studies, more frequent mood ratings could help determine the exact mechanism of taVNS on mood. Third, we provided evidence that acute taVNS boosts mood recovery after effort exertion, but we did not observe an association between taVNS-induced changes in mood and taVNS-induced changes in effort parameters. However, this might be due to differential modulatory effects of individual tasks since the exertion of cognitive tasks can interfere with one’s ability to regulate mood (Kohn et al., 2015), whereas physical effort can support mood regulation and improve mood (Berger & Motl, 2000). Thus, systematically evaluating taVNS effects on mood recovery after different task types would provide a more refined understanding of taVNS-induced mood recovery.

To summarize, mood is a crucial moderator of everyday engagement with tasks, yet it has not been shown if non-invasive taVNS can swiftly enhance mood. Here, we showed that acute taVNS improves positive mood recovery in a post stimulation phase, but not during the stimulation when the strain of the effortful tasks was most pronounced. Moreover, participants showed a stronger taVNS-induced increase of invigoration if they had lower baseline levels of positive mood. Collectively, our results highlight the importance of vagal afferent signaling in mood homeostasis highlighting a previously unanticipated role for taVNS in specifically boosting mood recovery after exertion. To conclude, these findings are highly relevant in light of the etiology of affective disorders that are characterized by prolonged periods of negative mood after stress or effort (McEwen, 1998). Therefore, our results might pave the way for integrating non-invasive taVNS as an adjunct therapy into the current affective treatment repertoire.

## Acknowledgement

We thank Monja P. Neuser, Caroline Burrasch, Franziska Müller, Sandra Neubert, Moritz Herkner, and Leonie Osthof for help with data acquisition. The study was supported by the University of Tübingen, Faculty of Medicine fortune grant #2453-0-0. NBK received support from the Daimler and Benz Foundation, grant 32-04/19, the Else Kröner-Fresenius Stiftung, grant 2017_A67, and the Durham-Tübingen Seedcorn fund, grant DFG ZUK 63.

## Author contributions

NBK was responsible for the study concept and design. MF collected data under supervision by NBK. NBK conceived the method and MF & NBK processed the data and performed the data analysis. MF, VT, WL, FK, AK, & NBK wrote the manuscript. All authors contributed to the interpretation of findings, provided critical revision of the manuscript for important intellectual content, and approved the final version for publication.

